# Difference in hoof conformation between shod and barefoot-managed hooves

**DOI:** 10.1101/2021.02.23.432452

**Authors:** J.N. De Klerk

## Abstract

**Reasons for performing the study:** Hoof conformation is linked to biomechanics of the hoof and injury occurrence. There is no scientific data if conformation differs between shod and barefoot-managed hooves.

**Objectives:** To investigate if and how shod and barefoot hooves differ in conformation.

**Study design:** Retrospective cohort study.

**Methods:** Standardised lateral, dorsopalmar/dorsoplantar and solar photographs of 98 shod and 69 barefoot-managed hooves were included. Thirty-six of the barefoot horses were farrier-managed, 33 were podiatrist-managed. Length and angular measurements produced nine conformation parameters; dorsopalmar/plantar balance, solar symmetry, toe angle, heel angle, heel/toe angle difference, heel width, splaying index, flaring index and frog size.

**Results:** Barefoot hooves showed significantly fewer underrun heels, steeper heel angles, wider heels, increased splaying, increased flaring and larger frog size compared to hooves of shod horses. Solar symmetry showed a significant difference in front hooves but not hind hooves (P=0.038, P=0.104) and toe angle was not significantly different (P=0.368, P=0.425). There was no significant difference in the conformation of barefoot farrier and podiatrist-managed front hooves, however there was a significant difference in the hind hooves: farrier-managed hooves had longer frogs and shorter toes, compared to podiatrist-managed hooves.

**Conclusions:** The significant differences in hoof conformation found should be considered when managing the individual horse, since hoof conformation affects loading of the internal structure of the hoof and hence influences aetiopathogenesis of hoof pathology.

## Introduction

Horses are shod for comfort over all surfaces, particularly hard or rough terrains, and to prevent mechanical damage to the hoof capsule, such as splitting, bruising, cracking and excessive wear [1]. Good shoeing technique ensures suitable mediolateral and dorsopalmar/dorsoplantar hoof balance and provides support for internal structures. Shoes can also influence load distribution, for example by fitting extensions to address imbalances [2]. Many acknowledge that horseshoes provide support and can prevent or improve many hoof-related pathologies.

Some parties challenge shoeing, believing that shoes are cruel [3] and cause discomfort and lameness. In the late 20^th^ century, a “natural” barefoot hoof management was introduced collaboratively between a veterinarian and a farrier [4]. However, these concepts were considered radical, as many horses endured hoof pain and sub-solar abscess formation during the post-shoe removal transition period. Now, less radical trimming programmes have been introduced, advocated by the Equine Podiatry Association and the Institute of Applied Equine Podiatry [5].

Trimming and shoeing influences hoof conformation which affects the loading of the structures within the hoof and higher up the limb. One degree change in solar angle induces a 4% change in the strain of the digital flexor tendon and hence pressure experienced by the navicular bone [6]. These findings were verified by the results of a study where hoof conformation differed significantly between lame horses suffering from different lesions in the hoof [7]. Hoof growth and management influences locomotor biomechanics [8, 9] and hoof conformation [10]. The latter study investigated how hoof conformation changed over a 16-month period once implementing standardised barefoot trimming. This work demonstrated that the toe increased in length, the heel angle steepened and that the frog contact area increased significantly [10]. These changes were considered beneficial, particularly the raised heel height. A 5% increase in heel angle results in significantly less stress and displacement of distal limb structures [11]. Furthermore, an increased contact-area of the frog aids the hoof’s haemodynamic mechanism by assisting healing, growth and energy dissipation [12, 13].

There is limited scientific evidence about how hoof management influences hoof conformation. This study compared hoof conformation of horses managed barefoot and shod and hypothesised there would be a significant difference in hoof conformation between the two groups. In particular:

1. Lateromedial asymmetry, e.g. splaying, is more common in barefoot-managed horses.
2. Dorsopalmar/plantar hoof imbalance such as under-run heels, boxy hooves and long toes are more common in shod horses
3. Barefoot horses have wider, larger frogs and larger width between heels than shod horses In the UK, trimming of hooves is not regulated and is not limited to farriers, but is routinely performed by podiatrists. Training of farriers and podiatrists differs and we hypothesise:
4. There will be a significant difference in hoof conformation between barefoot-managed hooves managed by farriers compared to podiatrists.

## Materials and Methods

### Horses

Ninety-eight shod and 69 barefoot hooves were included in the study from 46 horses. Twenty-four horses were geldings and 22 were mares. All horses included in the study were from a general leisure horse population, regularly exercised, but not in competition work or heavy training routines. The horses were between 5 and 22 years old with a mean±SD age of 12.1±4.8 years. Heights ranged from 12.3hh to 17.3hh. All horses were maintained exclusively either shod or barefoot for more than 18 months before the study and on a hoof management regimen of either shoeing with standard shoes or kept barefoot with trimming performed every 5-7 weeks in both groups. A farrier or podiatrist who had been working full time for no less than 2 years, had received certified training in their area of work, and had undergone continual professional development since being awarded their degree, performed the shoeing and hoof trimming. The horses were split into 2 groups; shod, barefoot and the latter was further subdivided in farrier-managed and podiatrist-managed horses (36 farrier-managed, 33 podiatrist-managed).

### Conformation Measurements

Frontal, lateral and solar views were obtained from each front and hind hoof within two days of shoeing or trimming using a digital camera (Samsung WB250): Frontal and lateral photographs were obtained centring on the centre of the hoof with the camera set at a 30cm distance away using of a custom-made rig. For the solar photograph, the camera was set to be 30cm away from the apex of the frog. Linear and angular measurements were performed in Image J (http://rsbweb.nih.gov/ij/links.html) (table 1). Additionally, circumference at the coronary band (cm) and circumference of the sole (cm) were measured with a measuring tape. Type of horse-build based on height, weight and stature (light-weight/medium-weight/heavy-weight), gender, age and exercise regimen was also recorded.

**TABLE 1.**
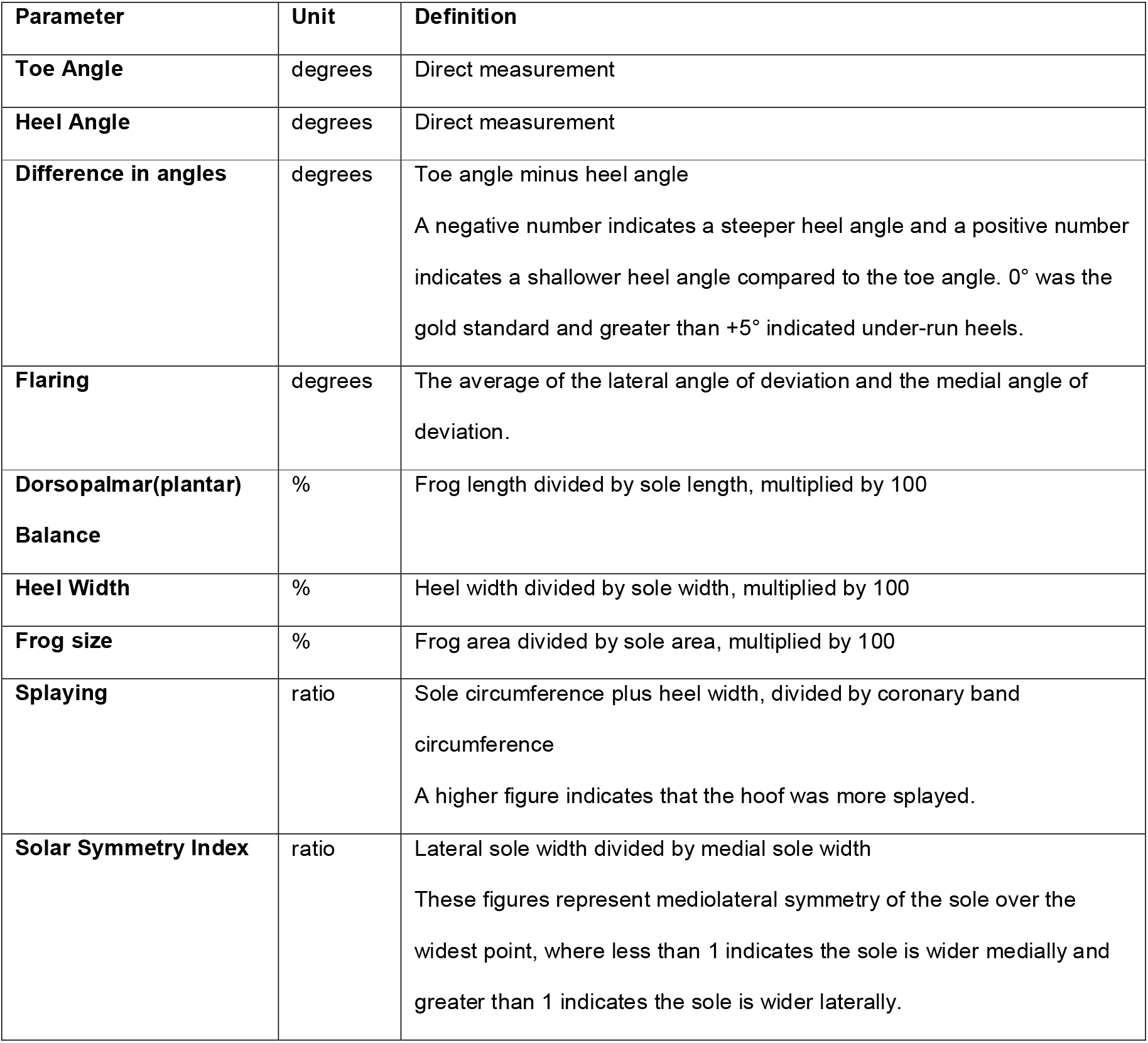
Conformation parameters included in this study

A repeatability assessment of the measurement technique was performed by measuring each parameter twice and calculating the limits of agreement [14, 15] (table 2).

**TABLE 2.**
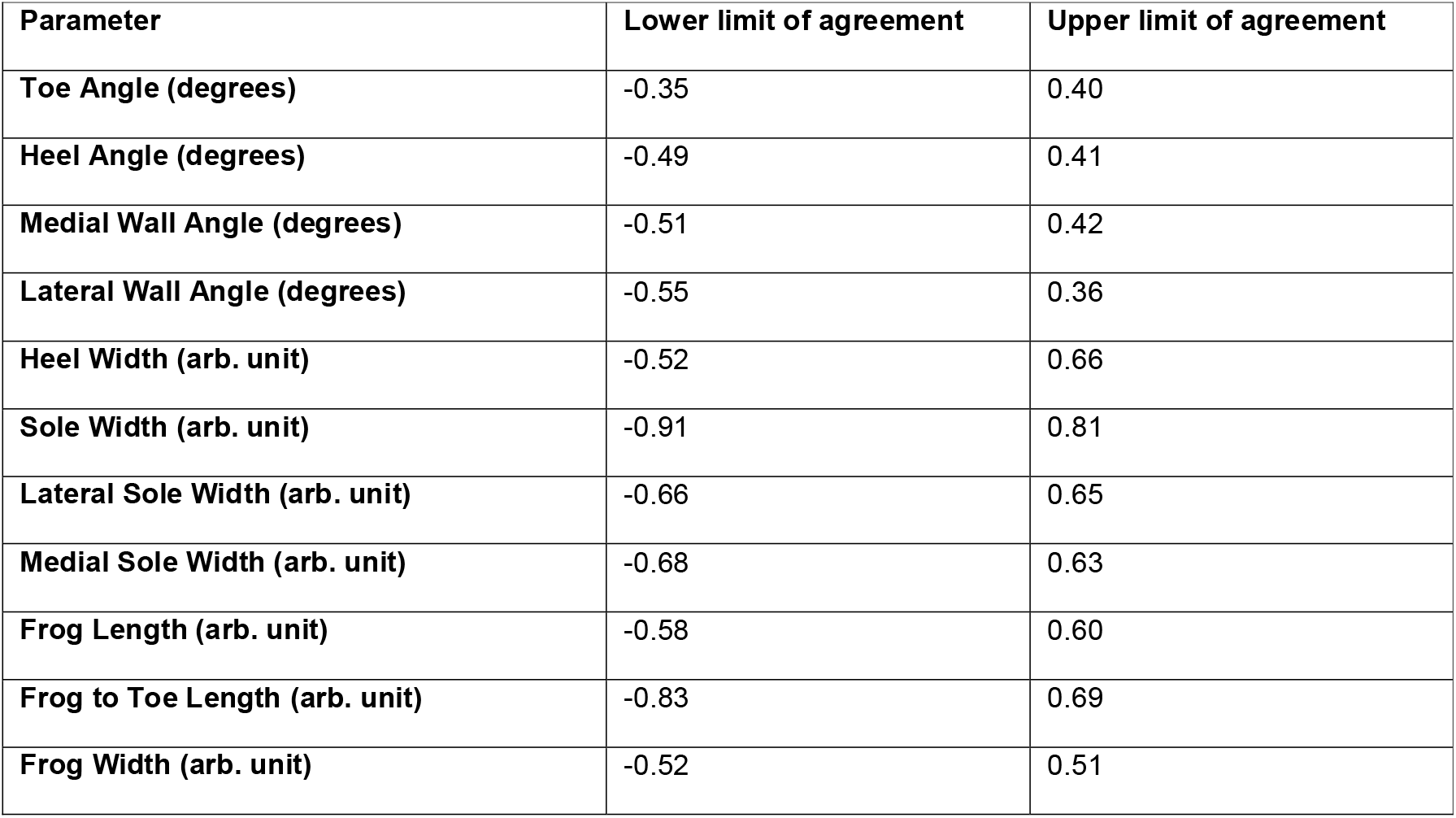
Reliability of conformation measurements: 95% limits of agreement for repeat measurements

### Data analysis

Data were tested for normality using the Shapiro-Wilk test and conformation parameters were compared between the shod and barefoot-managed hooves and between farrier and podiatrist-managed hooves using a T-test or Mann Whitney U test depending on the distribution of the individual conformation parameter. Horses were also grouped according to their stature and stature-type was also included in the analysis using a Kruskal-Wallis or One Way ANOVA test to assess if it was a confounding factor. When analysing stature-type, it was found that it was a confounding factor for dorsopalmar/dorsoplantar balance (P=0.004), heel angle (P=0.013), and frog size (P=0.001) and so it was ensured that the same ratio of light-weight, medium-weight and heavy-weight horses were included in each group for further statistical analysis. Left and right hooves were also analysed to ensure the values could be combined. For all parameters, there was no statistical significance, so left and right hooves were combined. All parameters were analysed in SPSS (IBM, version 22.0.0.1). P was set at 0.05.

## Results

### Comparison of conformation parameters of shod and barefoot-managed horses

Table 3 shows the hoof conformation parameters of the hooves measured in this study for shod and barefoot-managed hooves.

**TABLE 3.**
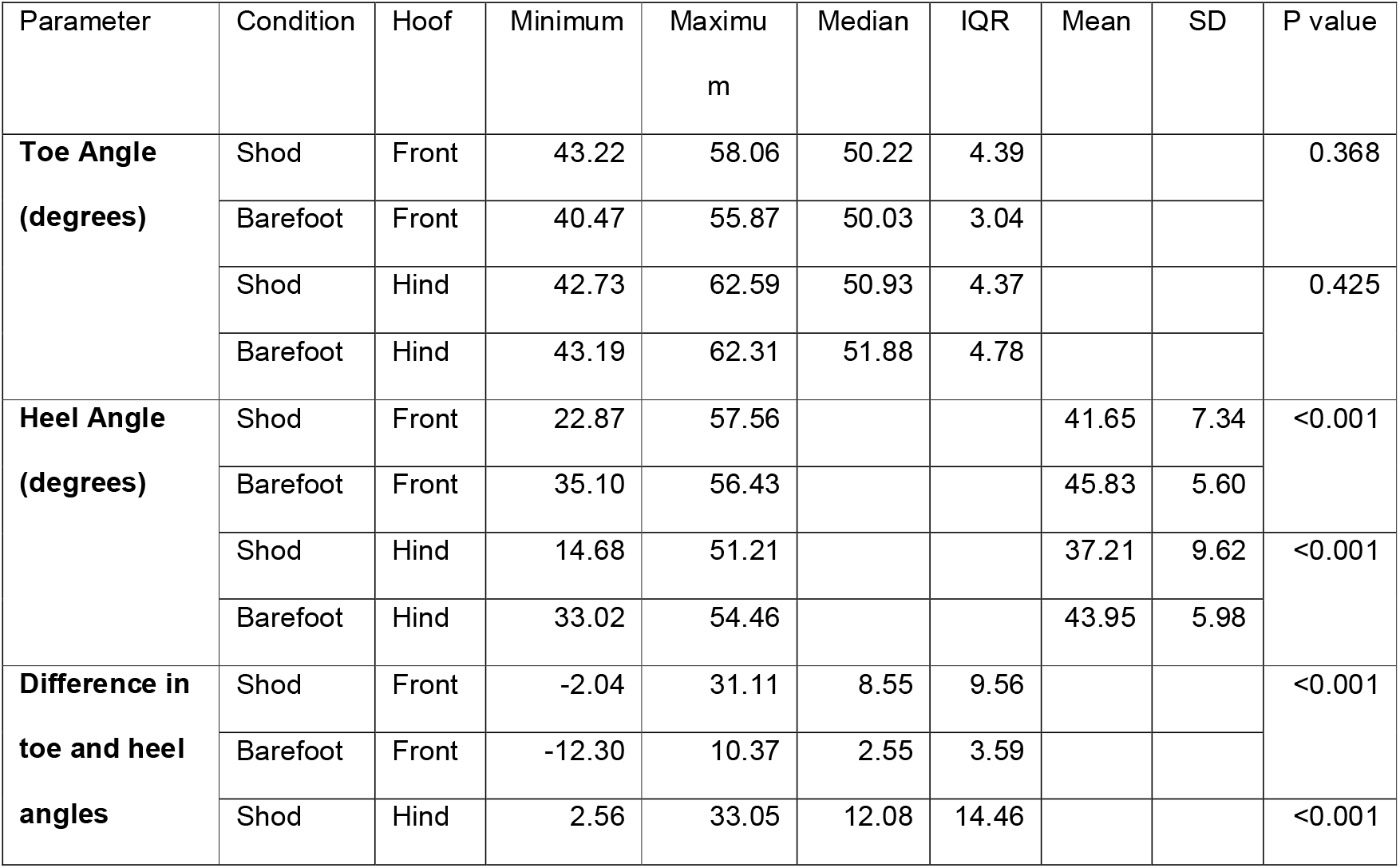
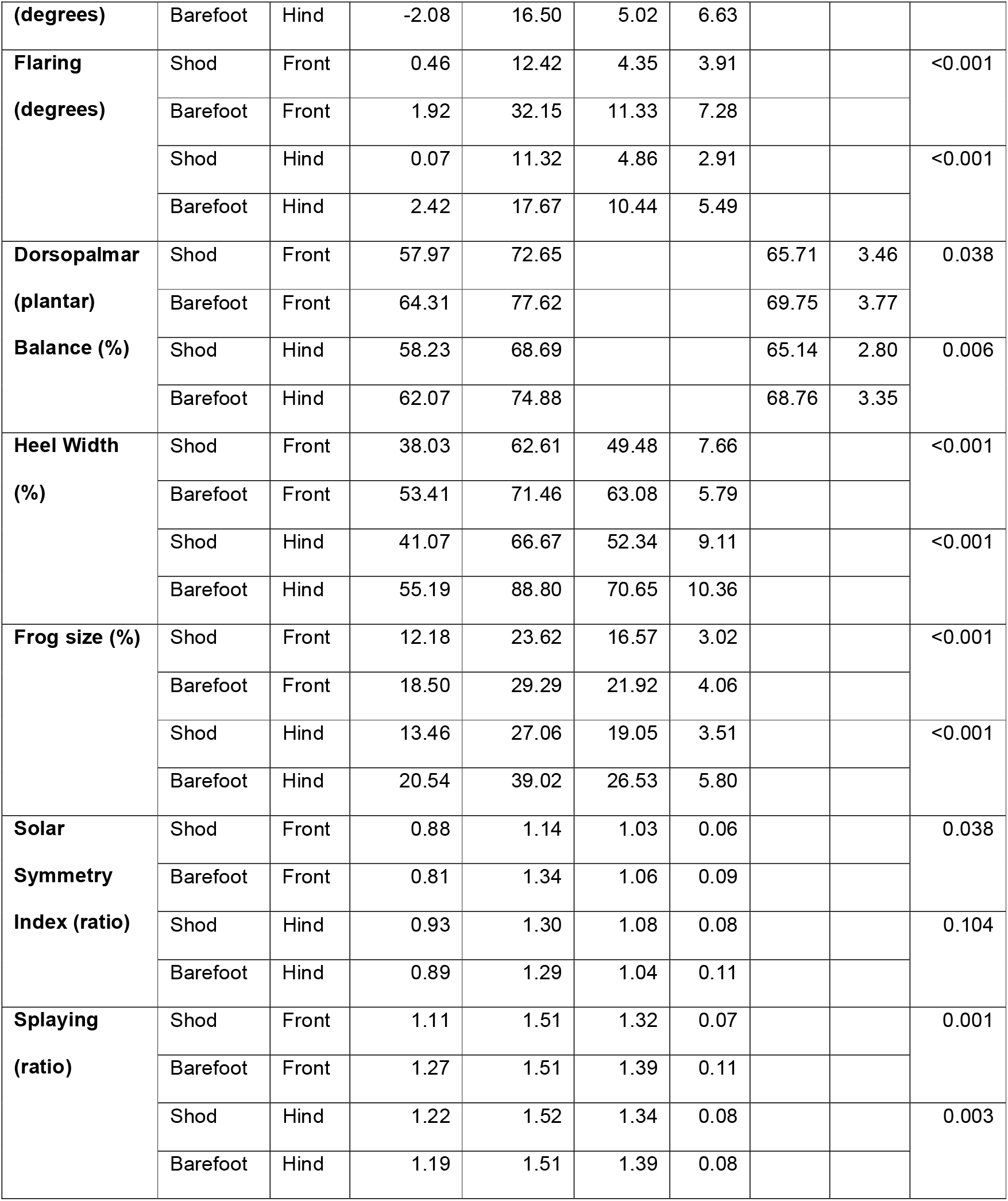
Conformation parameters for shod hooves and barefoot hooves for front and hind hooves. Mean and standard deviation (SD) are displayed for normally distributed parameters, median and interquartile range (IQR) are displayed for not-normally distributed parameters; P-value is for comparison between shod and barefoot-managed hooves.

The toe angle in front and hind hooves did not differ significantly between shod and barefoot-managed hooves. The heel angle however differed significantly between the two groups in both front and hind hooves. Shod hooves had a shallower heel angle compared to barefoot-managed hooves. In front hooves, the range of toe angle was much greater in shod hooves than in the barefoot hooves and the heel angle range was much greater for shod hooves in both, front and hind feet. There was a significant difference between the two groups in the difference in toe and heel angles in both front and hind hooves. Shod hooves had larger differences between toe and heel angles than barefoot hooves. There was a high prevalence of under-run heels, defined by a difference of more than 5 degrees [16], observed in both groups. 75.5% of hooves were under-run in the shod group and 40.5% were under-run in the barefoot group.

Barefoot-managed hooves showed a significantly greater heel width compared to shod hooves in front and hind. There was a significant difference in dorsopalmar/dorsoplantar balance between both groups for both front and hind hooves with barefoot hooves showing significantly shorter toes compared to shod hooves.

For both hoof management regimens, the hooves were on average wider laterally. In the front hooves, barefoot hooves were significantly more asymmetrical compared to shod hooves, but no significant difference in asymmetry was found between the two groups in the hind hooves.

There was a significant difference between hoof management regimens for splaying and flaring in both front and hind hooves with barefoot hooves being more splayed and flared than shod hooves.

There was a significant difference between hoof management regimens for frog area in both front and hind hooves with barefoot hooves having significantly larger frogs than shod hooves.

### Comparison of conformation parameters of barefoot hooves managed by a podiatrist and managed by a farrier

Table 4 shows the difference in hoof conformation parameters of barefoot hooves managed by a farrier compared to a podiatrist.

**TABLE 4.**
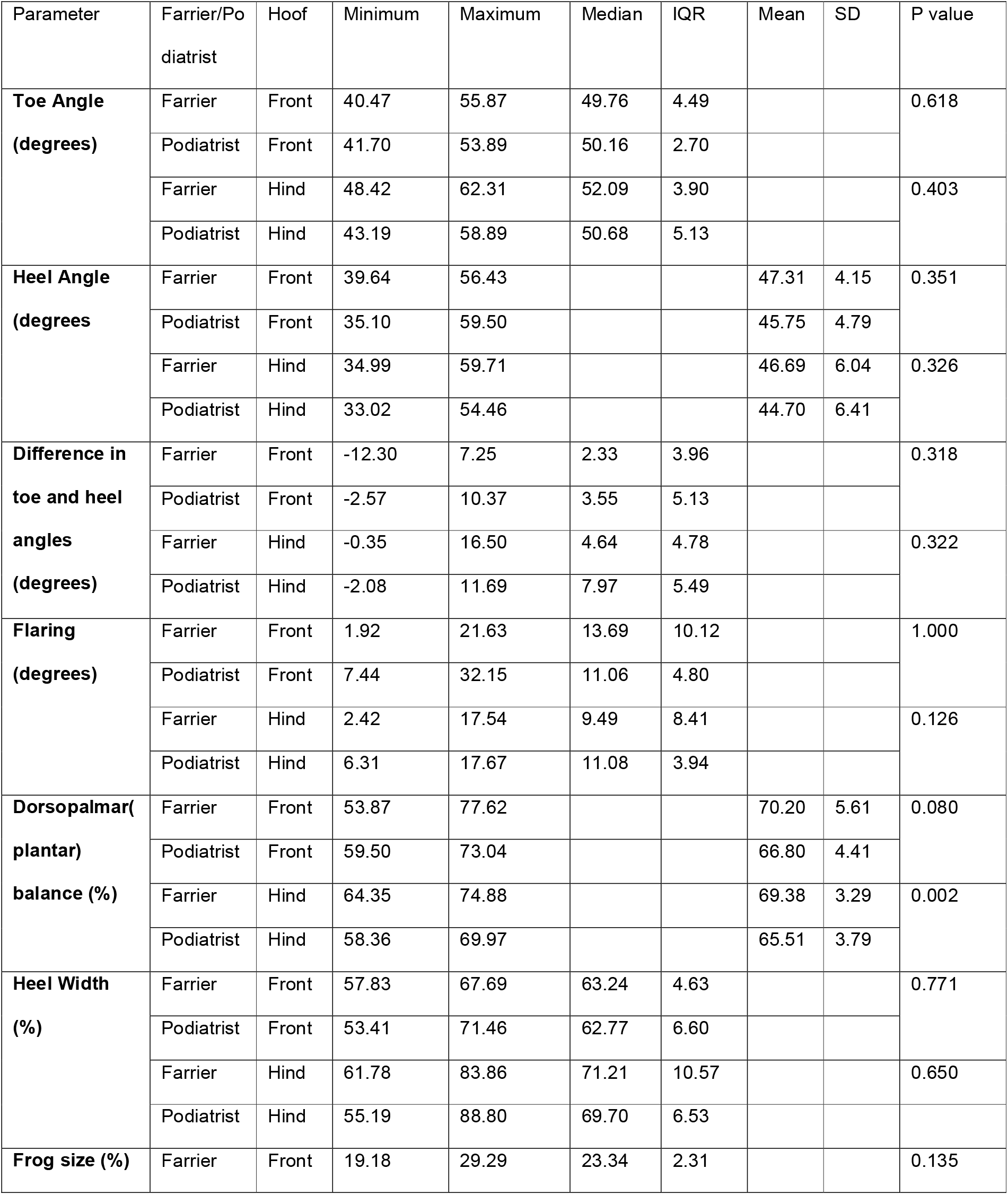
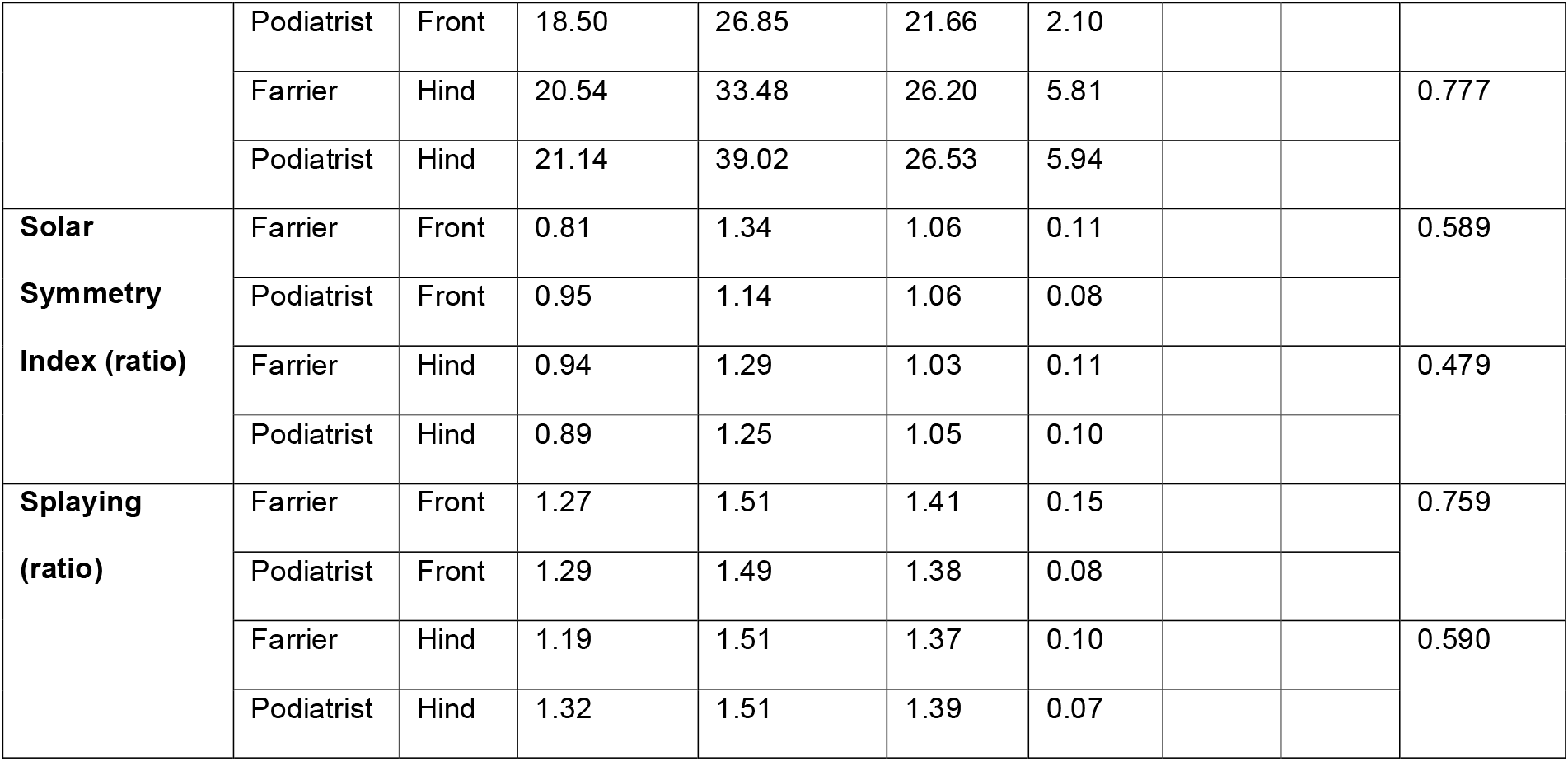
Conformation parameters for barefoot hooves managed by a farrier and barefoot hooves managed by a podiatrist for front and hind hooves. Mean and standard deviation (SD) are displayed for normally distributed parameters, median and interquartile range (IQR) are displayed for not-normally distributed parameters; P-value is for comparison between farrier and podiatrist-managed hooves.

There was no significant difference between farrier and podiatrist-managed front and hind hooves for heel angle, solar symmetry, toe angle, difference in toe and heel angles, heel width, splaying, flaring or frog size.

There was no significant difference between farrier and podiatrist-managed front hooves for dorsopalmar balance, however there was a significant difference in the hind hooves. Farrier-managed hooves had longer frogs and shorter toes, compared to podiatrist-managed hooves.

## Discussion

This study showed that some hoof conformation parameters differed significantly between shod and barefoot-managed hooves and between farrier and podiatrist-managed barefoot hooves.

In concordance to other recent studies on hoof conformation, it was found that hooves were wider laterally than medially in the forelimbs [17] and that hooves were more asymmetrical without shoes in the hindlimbs [18]. In agreement with previous findings there was a relationship between management regime and palmar angle, dorsal to palmar angle ratio, lateral heel height, sole length, sole length to width ratio, medial hoof width and solar symmetry [17]. Traditionally, in past literature, hoof angles were described to be around 45° in the forelimb and 55° in the hindlimb [1], which would suggest the results of this study were slightly steep in the forelimb and slightly shallow in the hindlimb, however this study’s results are consistent with other studies conducted in the past year where hoof angles were on average 50.3° in the forelimb [17] and 51.52°-52.16° in the hindlimb [18], suggesting a need to review hoof conformation of the modern domestic horse.

In the presented study there was no significant difference in toe angle between shod and barefoot-managed hooves, however the heel angle was significantly shallower in shod hooves compared to barefoot hooves. This may be due to the fact that these horses were more likely to be managed with shoes or it may be due to the fact that shoes increase the pressure on the horn in the heel area and over time lead to lowering of the heels. It has been shown previously that when hooves are managed without shoes, the distal half of the heels migrate palmarly/plantarly [10], thus increasing heel height. Lower heels are usually associated with a shallower solar angle of the distal phalanx and it has been shown that for each degree change in angle the strain and thus the stress on the navicular bone increases [6]. The opposite effect is seen when heel wedges are applied to increase heel angle, this results in a decrease of navicular bone stress in comparison to flat shoes. The same study showed that barefoot hooves showed a 14% reduction in the forces acting on the navicular bone, possibly due to the same mechanism as wedges [19]. Shallower heel angles may progress to under-run heels, which decrease the capability of the heels to deform under load thereby negatively impacting shock absorption and possibly also compromising the blood supply to the hoof [20]. This consequently may predispose to hoof pathologies. Hyperextension of the interphalangeal and fetlock joints also occurs with under-run heels, thereby increasing strain on the deep digital flexor tendon and the navicular bone [21].

Our study revealed that barefoot horses had more splayed and flared hooves than shod horses, suggesting that horseshoes might restrict multidirectional expansion of the hoof capsule. Shoes have been shown to decrease the deformation of the hoof wall during loading, by redistributing irregular strains without compromising hoof function, thereby protecting against flare [22]. Flare is undesirable in hooves, as it considerably weakens the hoof wall by bending of horn tubules [23].

Overall, all hooves in all groups were wider laterally. Previous studies have demonstrated that horses preferentially land laterally [17, 24] which corresponds to a more splayed, lower lateral and a steeper, higher medial hoof wall [25]. Slight asymmetry throughout all groups to the lateral side to a degree, is therefore considered normal. In the presented study there was only a significant difference in solar symmetry between hoof management regimens for front hooves, with barefoot hooves being more asymmetrical than shod hooves. The difference between the front and the hind hooves may be attributed to the uneven weight distribution of the horse with 58% of the weight being loaded on the forelimbs, and 42% loaded on the hindlimbs [26], as a greater downwards pressure on the hooves may increase lateral splay. Incorrect trimming, pain, or compensation for poor conformation may result in excessive lateral or medial landing and mediolateral imbalance [27]. Excessive asymmetry is important to address as it affects rotation, abduction and adduction of the distal interphalangeal joint [28], thereby potentially predisposing the distal limb to injuries relating to the articular surfaces and collateral ligaments [28,29], as well as causing chronic heel pain, sheared heels, metacarpophalangeal synovitis and side-bones [30]. The point at which asymmetry is considered excessive has not been quantified, however in this study a wider range of asymmetry was observed in barefoot hooves compared to shod hooves. The potential for barefoot-managed hooves to show more hoof asymmetry and hence for more uneven loading ought to be taken into account when dealing with those horses.

Heel width and frog size were both significantly associated with hoof management regimen. Barefoot hooves had wider heels and larger frogs than shod hooves. A wide heel and large frog is considered an advantageous feature of the hoof since it increases the load-bearing area, thus reducing stress. An increased area in the palmar/plantar aspect of the solar dermis could also potentially increase proprioception of the horse during locomotion, due to the high local concentration of Pacinian corpuscles [31]. It has been suggested that the frog hypertrophies when it becomes a weight-bearing structure [32]. Barefoot frogs have more stimuli for growth as they impact the ground, instead of being raised above it like in shod hooves [10]. This is advantageous for the distal limb as increased frog-ground contact increases blood flow, thereby aiding healing, energy dissipation and growth [10,12, 13]

There was a significant difference in hindlimb, but not frontlimb conformation between farrier-managed and podiatrist-managed barefoot hooves with the farrier-managed horses showing significantly shorter hooves and longer frogs in the hindlimbs. This may be due to different approaches to trimming hindfeet in particular based on the different training members of both groups undergo. No significant differences in the frontlimb were observed despite the proclaimed fundamental difference in approach by the two groups. Further studies on the detailed differences of the individual training programmes in relation to trimming are necessary to investigate this further.

## Acknowledgements

The author wishes to acknowledge Mr Neil Nevison for constructing a photographic rig to the author’s design and Ms Jenny Butler-Smith, Miss Emma Delport, Mrs Deborah Mead, Mrs Kirsty Stuart, Mrs Diane Goddard, Mrs Shirley Exall, Miss Eva Cox, Mrs Marianne Cox, Ms Sarah Mollett, Miss Lisa Housden, Miss Harriet Bradford, Mrs Auriol Thorne, Ms Bevni Lyons, Miss Emma Burston, Mr Wayne Upton, Mr Jack Hopson, Ms Teresa Mallia, Ms Teresa Watson, Miss Jo Ricey and Miss Natasha Crawley for aiding in the recruitment of horses for the study.

## Ethical animal research

Ethical approval was granted by the Royal Veterinary College on 19^th^ March 2013 (Reference Number 2013/R62).

---

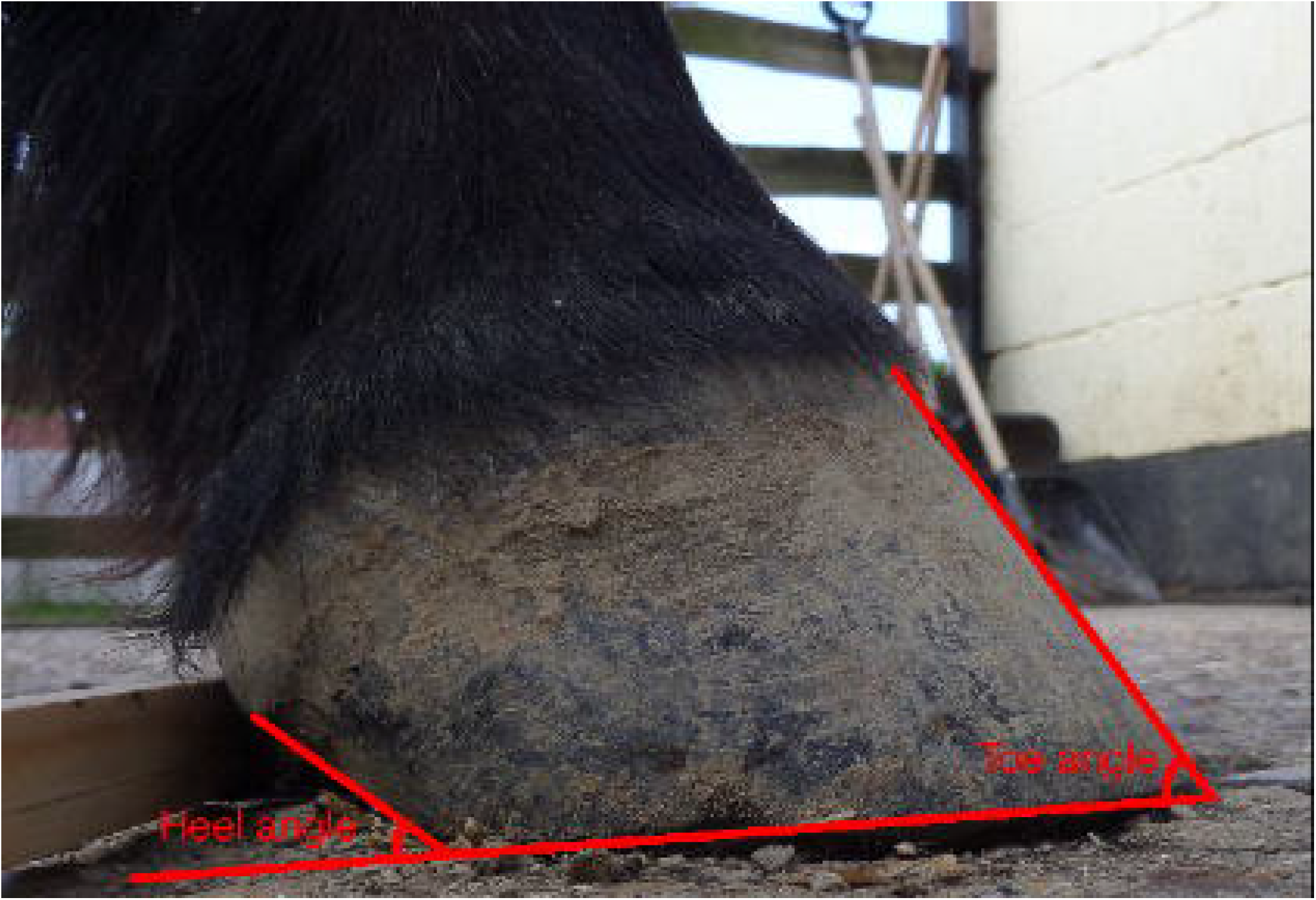

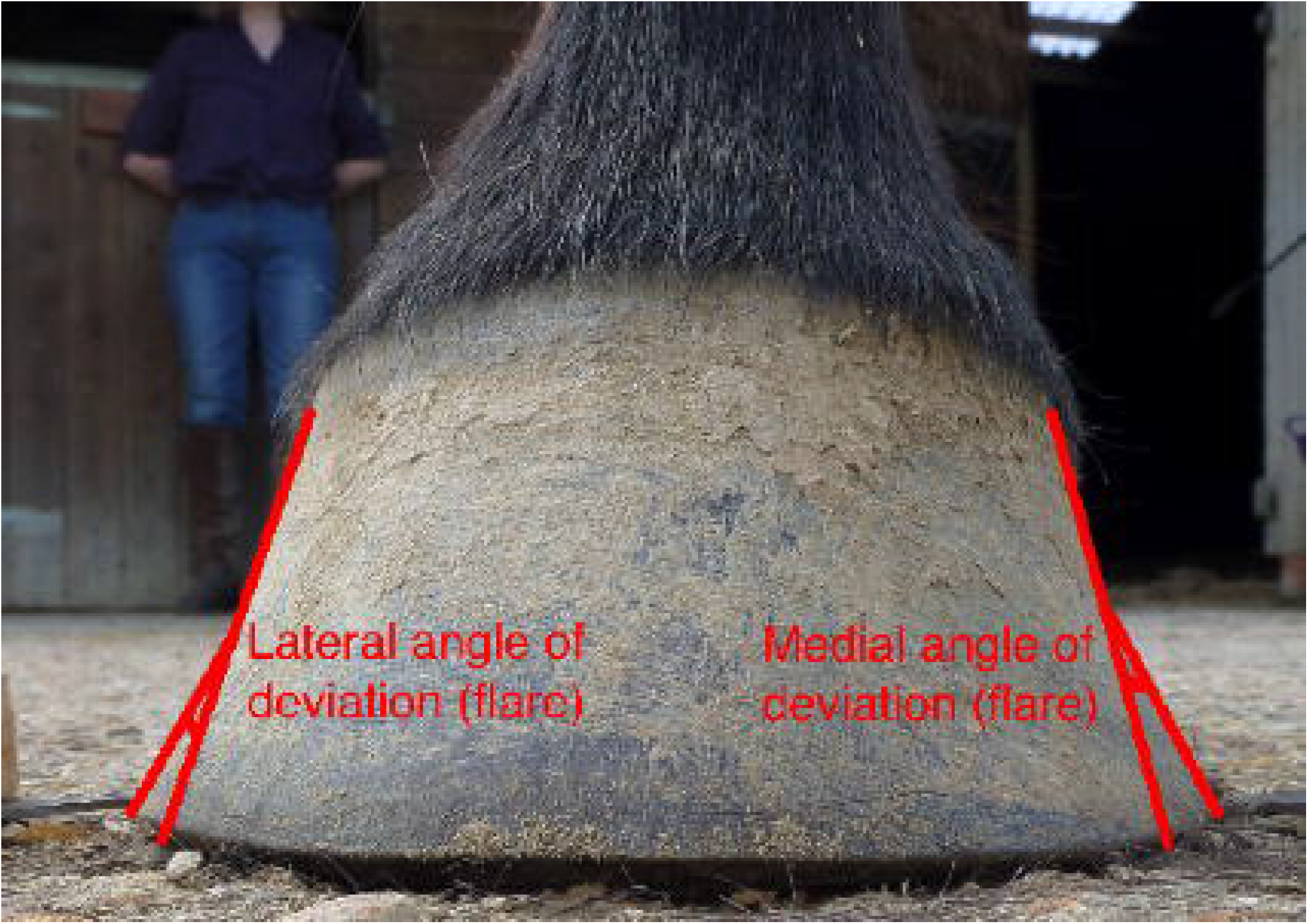

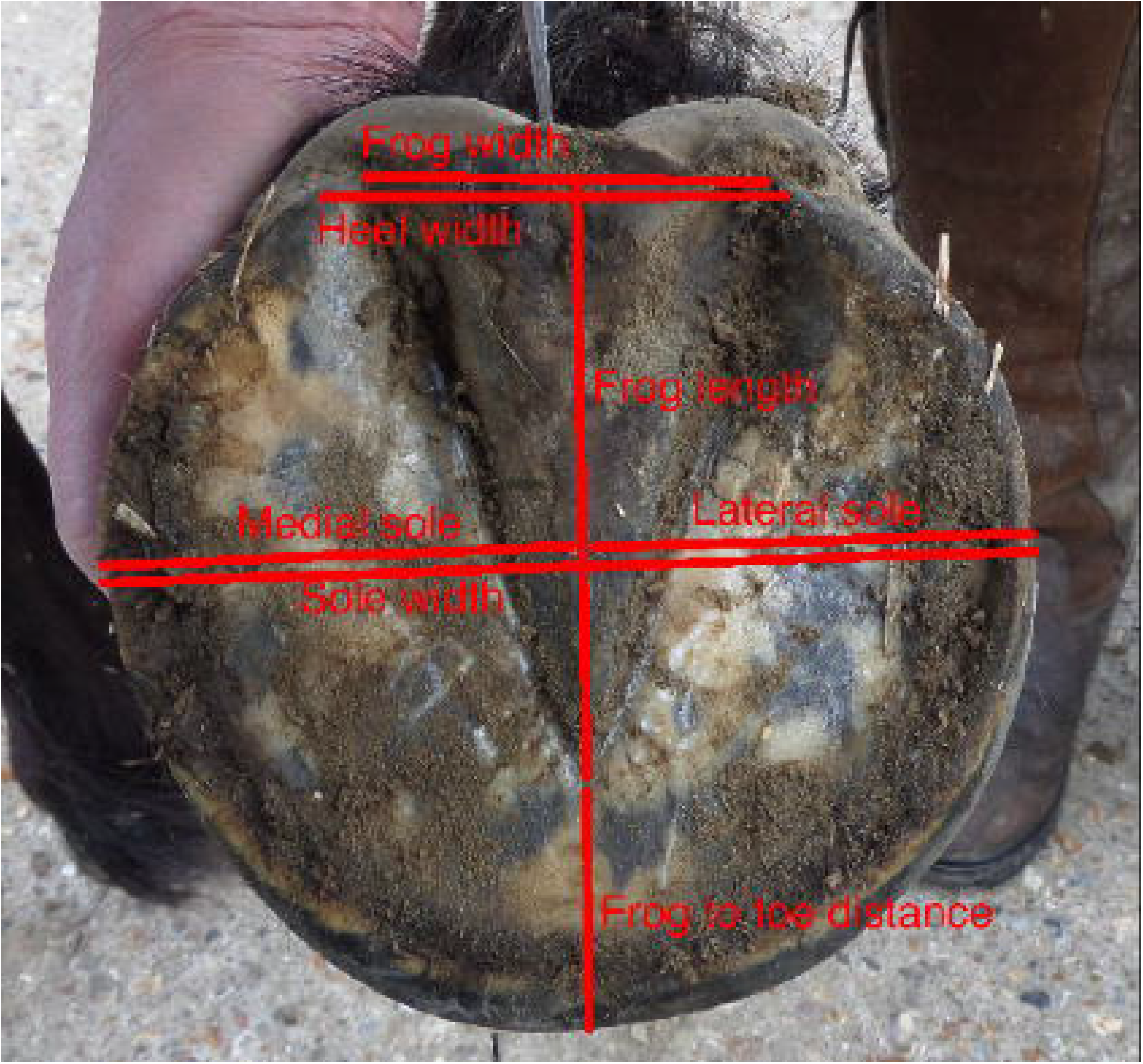

## Notes

### Competing Interest Statement

The authors have declared no competing interest.

