## Supplemental tables for "Difference in hoof conformation between shod and barefoot-managed hooves"

**TABLE 1 Conformation parameters included in this study**

| **Parameter** | **Unit** | **Definition** |
| --- | --- | --- |
| **Toe Angle** | degrees | Direct measurement |
| **Heel Angle** | degrees | Direct measurement |
| **Difference in angles** | degrees | Toe angle minus heel angle  A negative number indicates a steeper heel angle and a positive number indicates a shallower heel angle compared to the toe angle. 0° was the gold standard and greater than +5° indicated under-run heels. |
| **Flaring** | degrees | The average of the lateral angle of deviation and the medial angle of deviation. |
| **Dorsopalmar(plantar) Balance** | % | Frog length divided by sole length, multiplied by 100 |
| **Heel Width** | % | Heel width divided by sole width, multiplied by 100 |
| **Frog size** | % | Frog area divided by sole area, multiplied by 100 |
| **Splaying** | ratio | Sole circumference plus heel width, divided by coronary band circumference  A higher figure indicates that the hoof was more splayed. |
| **Solar Symmetry Index** | ratio | Lateral sole width divided by medial sole width  These figures represent mediolateral symmetry of the sole over the widest point, where less than 1 indicates the sole is wider medially and greater than 1 indicates the sole is wider laterally. |

**TABLE 2 Reliability of conformation measurements: 95% limits of agreement for repeat measurements**

| **Parameter** | **Lower limit of agreement** | **Upper limit of agreement** |
| --- | --- | --- |
| **Toe Angle (degrees)** | -0.35 | 0.40 |
| **Heel Angle (degrees)** | -0.49 | 0.41 |
| **Medial Wall Angle (degrees)** | -0.51 | 0.42 |
| **Lateral Wall Angle (degrees)** | -0.55 | 0.36 |
| **Heel Width (arb. unit)** | -0.52 | 0.66 |
| **Sole Width (arb. unit)** | -0.91 | 0.81 |
| **Lateral Sole Width (arb. unit)** | -0.66 | 0.65 |
| **Medial Sole Width (arb. unit)** | -0.68 | 0.63 |
| **Frog Length (arb. unit)** | -0.58 | 0.60 |
| **Frog to Toe Length (arb. unit)** | -0.83 | 0.69 |
| **Frog Width (arb. unit)** | -0.52 | 0.51 |

**TABLE 3 Conformation parameters for shod hooves and barefoot hooves for front and hind hooves. Mean and standard deviation (SD) are displayed for normally distributed parameters, median and interquartile range (IQR) are displayed for not-normally distributed parameters; P-value is for comparison between shod and barefoot-managed hooves.**

| Parameter | Condition | Hoof | Minimum | Maximum | Median | IQR | Mean | SD | P value |
| --- | --- | --- | --- | --- | --- | --- | --- | --- | --- |
| **Toe Angle (degrees)** | Shod | Front | 43.22 | 58.06 | 50.22 | 4.39 |  |  | 0.368 |
|  | Barefoot | Front | 40.47 | 55.87 | 50.03 | 3.04 |  |  |  |
|  | Shod | Hind | 42.73 | 62.59 | 50.93 | 4.37 |  |  | 0.425 |
|  | Barefoot | Hind | 43.19 | 62.31 | 51.88 | 4.78 |  |  |  |
| **Heel Angle (degrees)** | Shod | Front | 22.87 | 57.56 |  |  | 41.65 | 7.34 | <0.001 |
|  | Barefoot | Front | 35.10 | 56.43 |  |  | 45.83 | 5.60 |  |
|  | Shod | Hind | 14.68 | 51.21 |  |  | 37.21 | 9.62 | <0.001 |
|  | Barefoot | Hind | 33.02 | 54.46 |  |  | 43.95 | 5.98 |  |
| **Difference in toe and heel angles (degrees)** | Shod | Front | -2.04 | 31.11 | 8.55 | 9.56 |  |  | <0.001 |
|  | Barefoot | Front | -12.30 | 10.37 | 2.55 | 3.59 |  |  |  |
|  | Shod | Hind | 2.56 | 33.05 | 12.08 | 14.46 |  |  | <0.001 |
|  | Barefoot | Hind | -2.08 | 16.50 | 5.02 | 6.63 |  |  |  |
| **Flaring (degrees)** | Shod | Front | 0.46 | 12.42 | 4.35 | 3.91 |  |  | <0.001 |
|  | Barefoot | Front | 1.92 | 32.15 | 11.33 | 7.28 |  |  |  |
|  | Shod | Hind | 0.07 | 11.32 | 4.86 | 2.91 |  |  | <0.001 |
|  | Barefoot | Hind | 2.42 | 17.67 | 10.44 | 5.49 |  |  |  |
| **Dorsopalmar (plantar)**  **Balance (%)** | Shod | Front | 57.97 | 72.65 |  |  | 65.71 | 3.46 | 0.038 |
|  | Barefoot | Front | 64.31 | 77.62 |  |  | 69.75 | 3.77 |  |
|  | Shod | Hind | 58.23 | 68.69 |  |  | 65.14 | 2.80 | 0.006 |
|  | Barefoot | Hind | 62.07 | 74.88 |  |  | 68.76 | 3.35 |  |
| **Heel Width (%)** | Shod | Front | 38.03 | 62.61 | 49.48 | 7.66 |  |  | <0.001 |
|  | Barefoot | Front | 53.41 | 71.46 | 63.08 | 5.79 |  |  |  |
|  | Shod | Hind | 41.07 | 66.67 | 52.34 | 9.11 |  |  | <0.001 |
|  | Barefoot | Hind | 55.19 | 88.80 | 70.65 | 10.36 |  |  |  |
| **Frog size (%)** | Shod | Front | 12.18 | 23.62 | 16.57 | 3.02 |  |  | <0.001 |
|  | Barefoot | Front | 18.50 | 29.29 | 21.92 | 4.06 |  |  |  |
|  | Shod | Hind | 13.46 | 27.06 | 19.05 | 3.51 |  |  | <0.001 |
|  | Barefoot | Hind | 20.54 | 39.02 | 26.53 | 5.80 |  |  |  |
| **Solar Symmetry Index (ratio)** | Shod | Front | 0.88 | 1.14 | 1.03 | 0.06 |  |  | 0.038 |
|  | Barefoot | Front | 0.81 | 1.34 | 1.06 | 0.09 |  |  |  |
|  | Shod | Hind | 0.93 | 1.30 | 1.08 | 0.08 |  |  | 0.104 |
|  | Barefoot | Hind | 0.89 | 1.29 | 1.04 | 0.11 |  |  |  |
| **Splaying (ratio)** | Shod | Front | 1.11 | 1.51 | 1.32 | 0.07 |  |  | 0.001 |
|  | Barefoot | Front | 1.27 | 1.51 | 1.39 | 0.11 |  |  |  |
|  | Shod | Hind | 1.22 | 1.52 | 1.34 | 0.08 |  |  | 0.003 |
|  | Barefoot | Hind | 1.19 | 1.51 | 1.39 | 0.08 |  |  |  |

**TABLE 4 Conformation parameters for barefoot hooves managed by a farrier and barefoot hooves managed by a podiatrist for front and hind hooves. Mean and standard deviation (SD) are displayed for normally distributed parameters, median and interquartile range (IQR) are displayed for not-normally distributed parameters; P-value is for comparison between farrier and podiatrist-managed hooves.**

| Parameter | Farrier/Podiatrist | Hoof | Minimum | Maximum | Median | IQR | Mean | SD | P value |
| --- | --- | --- | --- | --- | --- | --- | --- | --- | --- |
| **Toe Angle (degrees)** | Farrier | Front | 40.47 | 55.87 | 49.76 | 4.49 |  |  | 0.618 |
|  | Podiatrist | Front | 41.70 | 53.89 | 50.16 | 2.70 |  |  |  |
|  | Farrier | Hind | 48.42 | 62.31 | 52.09 | 3.90 |  |  | 0.403 |
|  | Podiatrist | Hind | 43.19 | 58.89 | 50.68 | 5.13 |  |  |  |
| **Heel Angle (degrees** | Farrier | Front | 39.64 | 56.43 |  |  | 47.31 | 4.15 | 0.351 |
|  | Podiatrist | Front | 35.10 | 59.50 |  |  | 45.75 | 4.79 |  |
|  | Farrier | Hind | 34.99 | 59.71 |  |  | 46.69 | 6.04 | 0.326 |
|  | Podiatrist | Hind | 33.02 | 54.46 |  |  | 44.70 | 6.41 |  |
| **Difference in toe and heel angles (degrees)** | Farrier | Front | -12.30 | 7.25 | 2.33 | 3.96 |  |  | 0.318 |
|  | Podiatrist | Front | -2.57 | 10.37 | 3.55 | 5.13 |  |  |  |
|  | Farrier | Hind | -0.35 | 16.50 | 4.64 | 4.78 |  |  | 0.322 |
|  | Podiatrist | Hind | -2.08 | 11.69 | 7.97 | 5.49 |  |  |  |
| **Flaring (degrees)** | Farrier | Front | 1.92 | 21.63 | 13.69 | 10.12 |  |  | 1.000 |
|  | Podiatrist | Front | 7.44 | 32.15 | 11.06 | 4.80 |  |  |  |
|  | Farrier | Hind | 2.42 | 17.54 | 9.49 | 8.41 |  |  | 0.126 |
|  | Podiatrist | Hind | 6.31 | 17.67 | 11.08 | 3.94 |  |  |  |
| **Dorsopalmar(plantar) balance (%)** | Farrier | Front | 53.87 | 77.62 |  |  | 70.20 | 5.61 | 0.080 |
|  | Podiatrist | Front | 59.50 | 73.04 |  |  | 66.80 | 4.41 |  |
|  | Farrier | Hind | 64.35 | 74.88 |  |  | 69.38 | 3.29 | 0.002 |
|  | Podiatrist | Hind | 58.36 | 69.97 |  |  | 65.51 | 3.79 |  |
| **Heel Width (%)** | Farrier | Front | 57.83 | 67.69 | 63.24 | 4.63 |  |  | 0.771 |
|  | Podiatrist | Front | 53.41 | 71.46 | 62.77 | 6.60 |  |  |  |
|  | Farrier | Hind | 61.78 | 83.86 | 71.21 | 10.57 |  |  | 0.650 |
|  | Podiatrist | Hind | 55.19 | 88.80 | 69.70 | 6.53 |  |  |  |
| **Frog size (%)** | Farrier | Front | 19.18 | 29.29 | 23.34 | 2.31 |  |  | 0.135 |
|  | Podiatrist | Front | 18.50 | 26.85 | 21.66 | 2.10 |  |  |  |
|  | Farrier | Hind | 20.54 | 33.48 | 26.20 | 5.81 |  |  | 0.777 |
|  | Podiatrist | Hind | 21.14 | 39.02 | 26.53 | 5.94 |  |  |  |
| **Solar Symmetry Index (ratio)** | Farrier | Front | 0.81 | 1.34 | 1.06 | 0.11 |  |  | 0.589 |
|  | Podiatrist | Front | 0.95 | 1.14 | 1.06 | 0.08 |  |  |  |
|  | Farrier | Hind | 0.94 | 1.29 | 1.03 | 0.11 |  |  | 0.479 |
|  | Podiatrist | Hind | 0.89 | 1.25 | 1.05 | 0.10 |  |  |  |
| **Splaying (ratio)** | Farrier | Front | 1.27 | 1.51 | 1.41 | 0.15 |  |  | 0.759 |
|  | Podiatrist | Front | 1.29 | 1.49 | 1.38 | 0.08 |  |  |  |
|  | Farrier | Hind | 1.19 | 1.51 | 1.37 | 0.10 |  |  | 0.590 |
|  | Podiatrist | Hind | 1.32 | 1.51 | 1.39 | 0.07 |  |  |  |
